# BlenderPhotonics – an integrated open-source software environment for 3-D meshing and photon simulations in complex tissues

**DOI:** 10.1101/2022.01.12.476124

**Authors:** Yuxuang Zhang, Qianqian Fang

## Abstract

**Significance:** Rapid advances in biophotonics techniques require quantitative, model-based computational approaches to obtain functional and structural information from increasingly complex and multi-scaled anatomies. The lack of efficient tools to accurately model tissue structures and subsequently perform quantitative multi-physics modeling greatly impedes the clinical translation of these modalities.

**Aim:** While the mesh-based Monte Carlo (MMC) method expands our capabilities in simulating complex tissues by using tetrahedral meshes, the generation of such domains often requires specialized meshing tools such as Iso2Mesh. Creating a simplified and intuitive interface for tissue anatomical modeling and optical simulations is essential towards making these advanced modeling techniques broadly accessible to the user community.

**Approach:** We responded to the above challenge by combining the powerful, open-source 3-D modeling software, Blender, with state-of-the-art 3-D mesh generation and MC simulation tools, utilizing the interactive graphical user interface (GUI) in Blender as the front-end to allow users to create complex tissue mesh models, and subsequently launch MMC light simulations.

**Results:** Here we present a tutorial to our newly developed Python-based Blender add-on – BlenderPhotonics – to interface with Iso2Mesh and MMC, allowing users to create, configure and refine complex simulation domains and run hardware-accelerated 3-D light simulations with only a few clicks. We provide a comprehensive introduction to this new tool and walk readers through 5 examples, ranging from simple shapes to sophisticated realistic tissue models.

**Conclusion:** BlenderPhotonics is user-friendly and open-source, leveraging the vastly rich ecosystem of Blender. It wraps advanced modeling capabilities within an easy-to-use and interactive interface. The latest software can be downloaded at http://mcx.space/bp.

## 1 Introduction

Model-based computational techniques play an essential role in today’s medical imaging, giving rise to an array of emerging functional imaging modalities that offer more specific and accurate diagnostic information at much lower costs and higher levels of patient safety.^1, 2^ For example, functional tomographic imaging techniques based on computationally solving inverse models, such as diffuse optical tomography (DOT), microwave tomography (MWT),^3^ photo-acoustic tomography (PAI)^4^ etc, have made ample advances in the past few decades and are increasingly used in clinical care and research. Novel image reconstruction techniques based on numerical computation enhance image contrast and reduce noise, enabling clinicians to more easily discern early-stage diseases.^5^ Furthermore, the rise of artificial intelligence in recent years has demonstrated transformative effects on medical imaging and offers another excellent example on how computation can improve modern healthcare.^6^ To develop efficient model-based imaging data analysis pipelines, one must address several major hurdles, including 1) shape-modeling of complex tissue anatomies, 2) accurate shape discretization in terms of mesh generation or rasterization, and 3) efficient multi-physics solvers that can utilize such discretized domain to quantitaively solve the respective forward problems. An easy-to-use and fully open-source computational platform built for such purpose will be of great value to the research community.

In recent decades, near-infrared (NIR) based optical imaging methods have shown great potential in a number of applications, such as breast cancer diagnosis, neoadjuvant chemotherapy monitoring, and functional brain imaging. The use of non-ionizing radiation in optical imaging makes it possible for long-term monitoring; its relatively low-cost and portable instruments also permit broad access^7^ when compared to conventional imaging modalities. In addition, optical imaging provides functional information regarding the tissue’s physiological status by recovering tissue chromophores (such as oxy-/deoxy-haemoglobin, water, lipids etc). For example, functional infrared spectroscopy (fNIRS) – an imaging technique that uses near-infrared light to detect brain activations^8^ – has been rapidly adopted in neuroscience research and clinical applications. In fNIRS, the spatiotemporal variations of oxy-hemoglobin (HbO) and deoxy-hemoglobin (HbR) concentrations due to brain activity can be detected with high temporal resolution using specific wavelengths in an optical window, where the absorption spectra of HbO and HbR are distinct.^9, 10^

Optical imaging relies on quantitative and accurate forward models to account for the complex photon-tissue interactions,^11, 12^ which can be described by the radiative transfer equation (RTE). The RTE connects photon radiance with the optical properties in the medium, i.e., absorption coefficient (*µ*_*a*_), scattering coefficient (*µ*_*s*_), anisotropy (*g*), and refractive index (*n*). Unfortunately, the RTE can only be solved analytically in simple domains, such as infinite media, semi-infinite media or infinite slabs; it has to be computed numerically in complex or random media.^13, 14^ The diffusion equation (DE), an approximation to the RTE, is only valid in domains where scattering is dominant, i.e. *µ*_*s*_ ≫ *µ*_*a*_. One can solve the DE efficiently using numerical methods, such as the finite-element method (FEM), over a discretized domain in the form of a tetrahedral mesh. A number of FEM based DE solvers have been reported, including NIRFAST,^15^ TOAST++^16^ and Redbird.^17^ Although the DE can be solved much more quickly when compared to RTE solvers,^18^ it may yield erroneous solutions in several tissue types, including non- or low-scattering tissues such as cerebrospinal fluid (CSF) regions in the brain, lungs, or nasal cavities. In such cases, solving the RTE becomes necessary.^19^

The Monte Carlo method (MC), a stochastic solver to the RTE, is widely recognized as the gold-standard for solving the RTE.^20^ In an MC simulation, photons are simulated in packets, characterized by a “weight” initialized as 1 and updated as the photon traverses through the domain. Unlike the DE, MC simulations are based on repeated random samplings of the probability of photon absorption and scattering processes, thereby requiring the simulation of large numbers of photons in order to produce convergent results.. In recent years, the high speed of graphics processing units (GPUs) and the “embarrassingly parallelizable” nature of MC have allowed the cost of computation to drop dramatically, resulting in hundreds or even thousands-fold speed improvements compared to traditional MC methods.^21, 22^

Currently, several commercial optical design software packages support MC-based photon simulations, such as Zemax^®^(Zemax LLC, WA, USA), and TracePro^®^(Lambda Research, MA, USA). These commercial optical design tools usually have an interactive computer-aided design (CAD) based graphical user interface (GUI), easy-to-use parameter settings, and intuitive 3-D renderings of simulation results. These features have made such optical design software attractive among commercial users and research laboratories. However, commercial software often lacks advanced features such as GPU acceleration and cloud computing support; they also lack the most up-to-date MC simulation techniques such as shape-based and mesh-based MC methods.^11^ In addition, expensive licensing costs limit wide-spread use of these tools.

On the other hand, open-source MC simulators have seen tremendous growth over the last decade, offering superior simulation speed, advanced capabilities, and versatile algorithms when compared to their commercial counterparts. Mesh-based Monte Carlo (MMC) and voxel-based MC simulator Monte Carlo eXtreme (MCX), first reported in 2010 and 2009,^11, 23^ respectively, are two examples of advanced, open-source MC software packages that have attracted sizable user communities. Particularly, MMC utilizes tetrahedral meshes, similar to those used by an FEM DE solver, to simulate complex anatomical structures with high flexibility and excellent accuracy when compared to voxel-based domain representations. In several reported benchmarks,^23^ MMC outperforms voxel-based MC in accuracy when simulating curved boundaries, yet requires only a fractional memory footprint. Recently, MMC gained GPU-acceleration and also added support for a wide range of CPUs and GPUs via the OpenCL programming framework^24^ and CUDA.^25^ It is worth mentioning that MMC remains an actively developed open-source project that is constantly being updated, growing in both new features and accuracy. For example, wide-field MMC was added by Yao *et al*. in 2016,^26^ which is particularly important for supporting the active development of spatial frequency domain imaging (SFDI) techniques in recent years. In addition, implicit mesh-based Monte Carlo algorithm (iMMC), proposed by Yuan *et al*.,^27^ combines shape-based modeling and mesh-based anatomical models to enable simulations of extremely complex tissue structures such as dense vessel networks.

Regardless of the method used, whether it be MMC-based MC simulations or FEM-based DE solvers, a tetrahedral mesh-based anatomical model is typically required. It is important to note that, in order to obtain accurate results in either approach, high quality meshes must first be generated. Open-source three-dimensional (3-D) meshing tools such as Iso2Mesh,^28^ CGAL^29–31^ and Tetgen,^32^ while widely adopted and capable of creating complex, high-quality mesh models from 3-D medical images, are largely designed for used in a command-line interface. Therefore, less-experienced users often encounter a steep learning curve when adopting these meshing tools for their applications. In addition, manually editing meshes or interactively fine-tuning mesh features in these command-line oriented meshing tools can be quite challenging or impossible. On the other hand, developments in the field of 3-D modeling and animation, driven by the game and movie industries, promise easy-to-use and highly interactive shape-based modeling software tools. This results in widely disseminated open-source 3-D modeling software, such as Blender,^33^ and commercial tools, such as Cinema-4D^®^(C4D, Maxon Computer GMBH, Germany) and Maya^®^(Autodesk, CA, USA). Most of these 3-D modeling tools provide visual interfaces and comprehensive model editing capabilities that are missing from typical meshing tools.

We would like to highlight Blender because it is a free and open-source 3-D modeling suite, widely used in the field of animation and digital media creation. It supports a comprehensive 3-D modeling pipeline: creation, animation, simulation, rendering and motion tracking. In 2020 alone, Blender was downloaded more than 14 million times. Supported by a large and active community, Blender greatly reduces the barrier of entry for beginners^33^ in 3-D modeling. To accommodate advanced programming needs, a Python application programming interface (API) published by Blender – `bpy` – is available for users to programmatically control Blender or develop add-ons. Over the past several years, Blender-based software packages for biomedical applications have been developed. Examples include BioBlender and ePMV.^34, 35^ The majority of these projects specifically focus on the processing and rendering of object/shape surfaces; therefore, a triangular surface mesh is typically used in their processing pipelines. These tools typically do not offer the capability to tessellate the interior space of objects, which is the essential task of 3-D mesh generation and mesh-based light transport simulations. A noticeable gap is presented between the surface-oriented 3-D modeling software and the need to discretize and quantitatively simulate the interior space bounded by these surfaces. Therefore, creating an interface between Blender and 3-D mesh generators and multi-physics simulators could readily transform Blender into a powerful 3-D quantitative simulation platform and benefit an array of computational imaging domains.

There are several Blender-based add-ons that implement 3-D voxelated volumetric data rendering, for example, OrtogOnBlender,^36^ which can visualize Digital Imaging and Communications in Medicine (DICOM) files in Blender and extract surface meshes from the DICOM image stack. However, these add-ons do not support tetrahedral mesh generation and thus can not be directly used for subsequent model-based analyses. In addition, the lack of fine-grained mesh quality control can also create challenges towards performing any quantitative modeling beyond rendering. In this work, Iso2Mesh, an open source MATLAB/GNU Octave-compatible mesh generator, is used to generate high-quality tetrahedral meshes ^28, 37^ using the initial surface models created by Blender. It is worth highlighting that GNU Octave is a free and open-source high-level numerical analysis platform that is largely compatible with MATLAB. Octave also provides a Python API named `oct2py` to interface with Python. Using the combination of `oct2py` and `bpy`, one can efficiently bridge between Blender and Octave using Python as the “glue” language. The goal of this work is to develop an interactive interface between MMC/Iso2Mesh and Blender. This interface needs to have the ability to interactively create and visualize the model as well as automating the mesh generation and execution of mesh-based MC photon simulations via easy-to-use settings.

Here we present a tutorial to introduce to the community an open-source Blender add-on – Blender-Photonics – for performing advanced tissue-optics MC simulations. It combines the visual modeling capabilities of Blender,^33^ the meshing capabilities of Iso2Mesh,^28^ and the light simulation capabilities of MMC^23, 24^ to allow users to visualize and interactively edit the model while also configure light simulation in an intuitive interface. In addition, we show examples on how to create realistic tissue models, including creating highly complex human hairs and rough surfaces, using this tool. BlenderPhotonics is written in the Python and Octave languages and has a modular file structure. Overall, BlenderPhotonics makes the optical simulation pipeline much easier to use and opens the doors for creating highly sophisticated tissue models for future studies.

In the following sections, we first introduce the overall workflow of BlenderPhotonics, followed by the implementation details of each processing step. In Section 3, we demonstrate several showcases on 3-D light simulations from each of the 3 supported input data types, including 1) constructive solid geometry (CSG) inputs using 3-D shape primitives, 2) triangular surface models and 3) 3-D medical image volumes. Furthermore, we demonstrate the possibility of creating life-like complex tissue models and perform quantitative analyses in two advanced examples, showing users 1) how to simulate skins of different roughness levels using rough-surface creation, and 2) how to simulate complex human hairs using the hair/particle systems in Blender. Finally, we summarize the strengths and limitations of the software and discuss the plan for future extensions of this platform.

## 2 BlenderPhotonics Interface Design and Key Features

The four major functions of BlenderPhotonics include 1) creating tetrahedral meshes from 3-D objects (*Blender2Mesh*), 2) importing and processing externally created surface meshes (*Surf2Mesh*), 3) creating surface and tetrahedral meshes from 3-D volumetric images and arrays (*Volume2Mesh*), and 4) executing mesh-based MC light transport simulations and rendering results (*Multiphysics Simulations*). Correspondingly, we design this add-on in a modular fashion. Although these four tasks are logically sequential, each task can work independently as long as the appropriate input is provided. One can selectively run only one of the tasks or the entire pipeline, starting from shape modeling to light simulation.

In the next few subsections, we will discuss the processing pipelines for various user input data types. In order to unambiguously describe the intermediate mesh data generated through the pipeline, here we use “surface mesh” [Fig. 1(a)] to refer to a shell of triangular (known as “tris” in Blender), quadrilateral (known as “quads” in Blender), polygonal (known as “N-gons” in Blender) patches, or parametric 2-D manifolds that only discretizes the surface shape of the domain regions; we use “volume mesh” [Fig. 1(b)], to describe a tetrahedral volumetric mesh that discretizes both the surface as well as the interior space of the 3-D domain; we also use the word “regional mesh” [Figs. 1(c)-1(d)] to refer to sub-regions of the volume mesh that are tagged with unique labels to represent tissue-specific regions. Most 3-D shape primitives supported in Blender, such as cubes, spheres and cylinders, are defined as a surface mesh internally or can be converted to a surface mesh. Both the volume mesh and regional mesh are tetrahedral mesh models containing internal nodes, created by Iso2Mesh/Tetgen^32^ using surface meshes as inputs. When the input surface mesh contains multiple objects or surface compartments (i.e. multiple enclosed regions), the tetrahedral mesh elements inside each compartment are assigned a unique label, typically an integer number, and the tetrahedral mesh of each label is a regional mesh.

**Fig 1.**
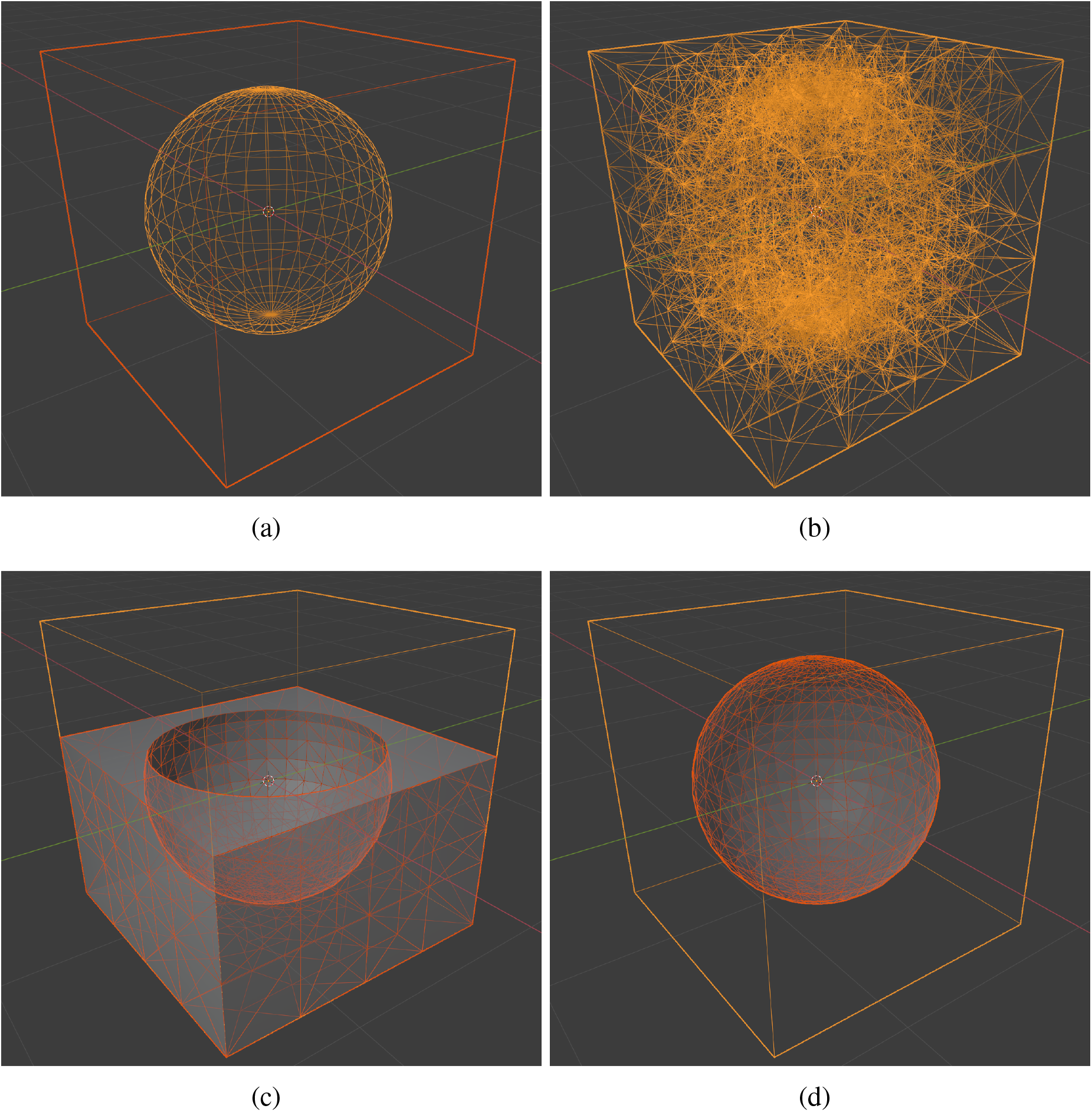
Different types of shape constructs used in Blender and BlenderPhotonics: (a) Blender objects (surface mesh) made of quadrilateral faces (“quads”) or polygons (“N-gons”), (b) volumetric tetrahedral mesh, (c) regional mesh of the cube excluding the sphere region (cropped), and (d) regional mesh of the center sphere.

### 2.1 Overall workflow

The overall workflow of this add-on is illustrated in Fig. 2. BlenderPhotonics contains three key steps. The first step is the construction of a surface mesh that is made of one or multiple objects. Users can create such surface models by using the built-in shape objects provided by Blender or importing from user-defined data files as an input. The second step is volume mesh generation and region labeling. The software exports the vertices and faces of the surface mesh generated from the first step and calls Iso2Mesh/Tetgen to populate tetrahedra to fill the enclosed compartments of the surface mesh – each region is uniquely numbered. To permit mesh density and quality fine tuning, a simple dialog is provided to allow the adjusting of meshing parameters interactively, followed by regeneration of the volume mesh. The third step is mesh-based MC light simulation using the volume or regional meshes generated from the second step. The optical properties of each tissue region and the light source parameters are configured visually via the “custom property” in Blender and an MMC compatible configuration data structure is generated, and is used as the input to launch the MMC simulation.The final light fluence distribution generated from such simulations, defined as floating-point fluence-rate values across the volume mesh, is then read in Blender and rendered for interactive visualization.

**Fig 2.**
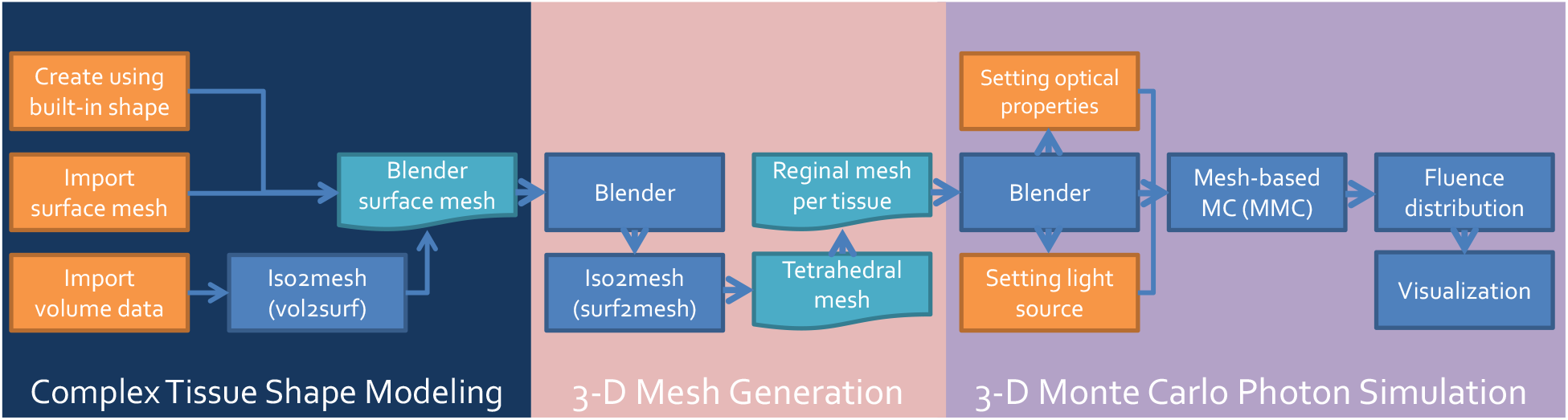
Overall workflow diagram of BlenderPhotonics.

The modular software structure shown in Fig. 2 makes it effortless for future extensions. If desired, one can potentially replace the MMC simulation module by another mesh-based MC simulator, such as FullMonteCUDA.^25^ Only lightweight code changes are required, including adding additional “custom properties” in Blender, exporting these additional user inputs in the format acceptable by the target simulator, and importing the output data back to Blender for rendering.

### 2.2 Installation

The installation of BlenderPhotonics requires setting up 6 software components: 1) Blender v2.8 or newer versions, 2) GNU Octave 4.2 or newer versions, 3) `oct2py` Python module (for interfacing Blender with GNU Octave), 4) `jdata` Python module (for reading/writing data exchange files^38^), 5) Iso2Mesh toolbox (for 3-D mesh generation), 6) MMCLAB toolbox (for photon MC simulation). Two optional components can be installed, including 1) ZMat toolbox (for reading/writing internally compressed mesh files) and 2) `bjdata` Python module (for reading binary JavaScript Object Notation (JSON) based^38^ data files). All required software components are open-source, and are widely available. The step-by-step installation instructions can be found in the “Installation” section of the README file on the Github repository.^39^

In our BlenderPhotonics package,^39^ we also provide a fully automated installation shell-script for a Debian/Ubuntu based Linux system. By simply replacing the `apt` command to `yum` or `port`, the attached installation script can be used to install BlenderPhotonics on Fedora Linux and Mac OS, respectively. For Windows users, although running the above shell script is possible if one has preinstalled Cygwin64, the `apt` command must be replaced because it is not supported on Windows. A Windows user may use `winget install -e --id BlenderFoundation.Blender`, `winget install -e --id GNU.Octave`, and `winget install -e --id coti.mcxstudio` in an administrator command window to install the needed software components, followed by installing the Python modules and Octave toolboxes manually.

Once all the software components are successfully installed, one should be able to start Blender, click on the menu *“Edit Preferences”*, choose *“Add-ons”* in the Blender Preferences window, and search for “BlenderPhotonics”, the downloaded add-on should be listed (or use the *“Install”* to browse locally downloaded zip package). To enable BlenderPhotonics, one should simply click on the check-box until a check-mark is shown. Although BlenderPhotonics has been tested on Blender 2.83, 2.92 and 3.0, the later versions are recommended because it supports an “exact” shape intersection solver starting from version 2.9. Aside from supporting GNU Octave, BlenderPhotonics also allows users to run Iso2Mesh and MMC via MATLAB (MathWorks, MA, USA) in the backend if one has installed the proprietary `matlab.engine` Python interface. One can choose between these two backends by a simple toggle-button at the top of the BlenderPhotonics interface (see Fig. 3).

**Fig 3.**
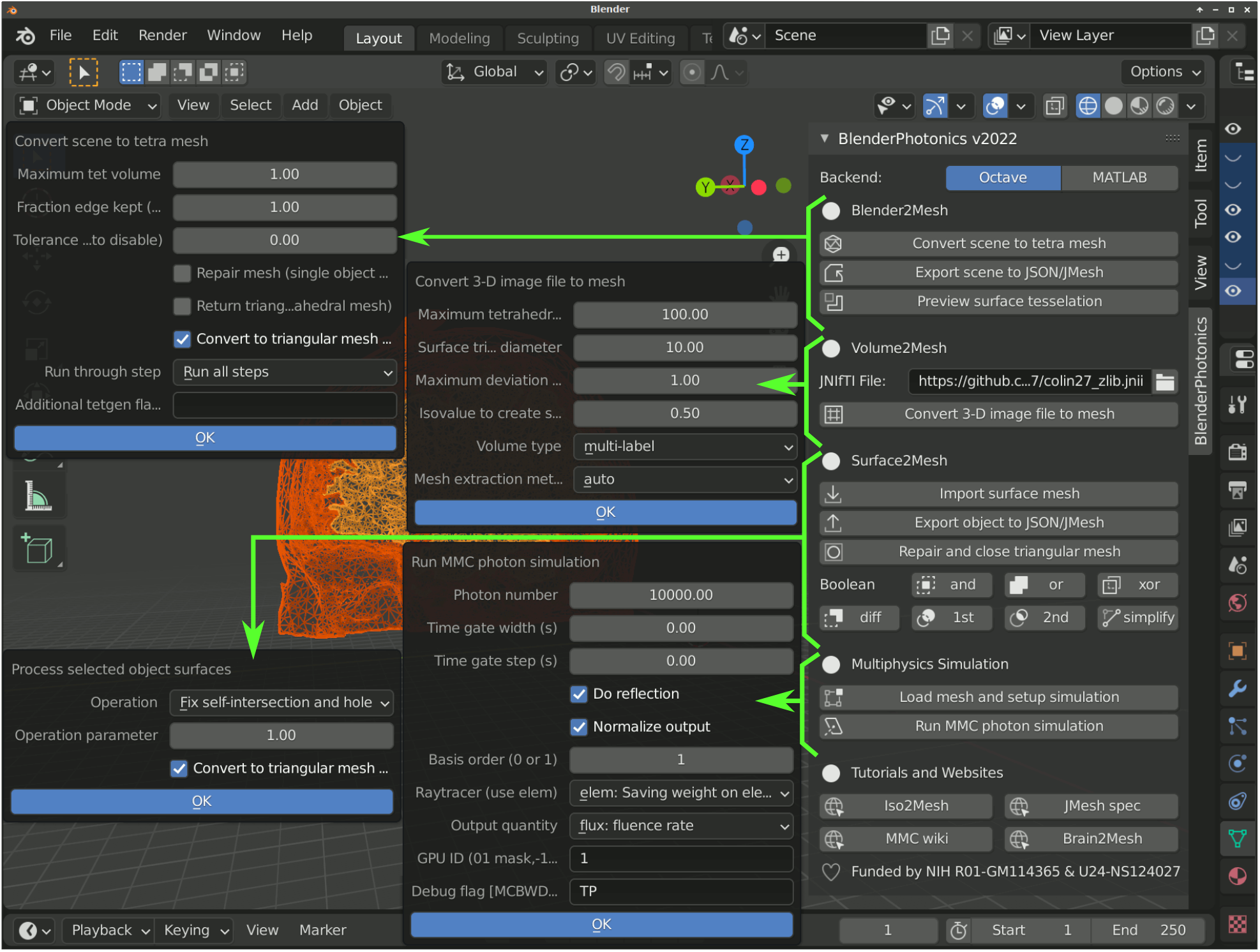
BlenderPhotonics main interface and parameter dialogs (pointed by arrows) for each core feature.

### 2.3 GUI design

Ease-of-use is one of the key design requirements of BlenderPhotonics. To allow novice users to conveniently access all the key functions of BlenderPhotonics, we created a simple function panel using Blender’s plug-in interface, see Fig. 3. In addition, it is desirable that a user is able to individually run any one of the functions. The independence of the task stages makes transferring or restarting portion of the processing pipeline possible – for example, using a portable computer to create the mesh model and then run GPU-based simulations on a dedicated server. The main interface allows to browse volume data files, create volumetric mesh, import regional mesh, and execute MMC simulation with only a single click.

### 2.4 Data inputs and model creation

BlenderPhotonics can create complex surface models (i.e. Step 1 in Fig. 2) based on various forms of user inputs. Overall, there are three types of inputs that can be used for model creation. First, a user can directly add one or multiple built-in shape objects, such as cubes and spheres, using Blender and create a complex domain via Blender’s vastly rich built-in tools, such as Boolean modifier, bisect, and knife tools etc, and then create triangular/tetrahedral meshes via the “*Blender2Mesh*” module in BlenderPhotonics (see Fig. 3). Secondly, a user can import a triangular surface mesh pre-generated by other tools, handled by the *“Surface2Mesh”* module in BlenderPhotonics. Thirdly, a 3-D voxelated image, in the form of a Neuroimaging Informatics Technology Initiative (NIfTI),^40^ JData^38^/JNIfTI^41^ or a MATLAB/Octave .mat file, can be imported to create an image-derived surface mesh as supported in the “*Volume2Mesh*” module in Blender-Photonics. Regardless of the input data type, the result of this step is a surface mesh defined by two data structures – vertices and faces. The vertices of the surface mesh is a floating-point array of dimensions *N*_*n*_ *×*3, with the 3 columns representing the *x/y/z* coordinates, respectively, of a vertex in the surface mesh, where *N*_*n*_ is the total number of nodes. The faces of the surface mesh are defined as a 2-D integer array of size *N*_*e*_ *×*3, with each row containing 3 integer indices of the 3 nodes that forms the triangular patch, where *N*_*e*_ is the number of triangles. Only closed (i.e. watertight) surface mesh models are accepted because it is required for the subsequent tetrahedral mesh generation and MC simulation. A user should also pay attention to the complexity, measured by the number of nodes and triangles, of the surface model. Obviously, exceedingly dense surface meshes can lead to very dense volumetric mesh, resulting in excessively long meshing time and subsequent MC simulation run-times. A best practice is to create the surface mesh with minimal number of nodes/triangles without losing the accuracy of the domain boundaries.

Blender is a vastly versatile environment for 3-D shape modeling and domain creation, and how to use most of the functionalities of Blender for complex shape creation is beyond the scope of this tutorial. Readers should browse the large number of tutorials created by Blender’s developers and user community to learn how to effectively build complex 3-D models using Blender. It is important to note that Blender separates built-in objects into (surface) mesh and non-mesh types, where the former has well-defined node coordinates and face node indices, and the latter may only have implicit surface definitions without explicitly defining surfaces or vertices. In many cases, non-mesh objects, such as a “metaball” object, can be converted to a mesh object by Blender during export. Other advanced built-in object types have not been tested.

A variety of surface mesh files can be imported to Blender. Blender supports most of the major 3-D model files such as OBJ geometry (.obj), STereoLithography (.stl), Filmbox (.fbx) formats, among others. By using BlenderPhotonics’s *“Surface2Mesh”* module and Iso2Mesh, we have also added support to import triangular surfaces stored in JSON/text-JMesh (.json or .jmsh), UBJSON/binary JMesh^42^ (.bmsh), object file (.off), Infria MEDIT (.medit) formats. In addition, Blender, and subsequently BlenderPhotonics, can have extended file format support when a user installs appropriate add-ons to read/write a customized data format. For external model files, most models defined in these inputs contain the required vertex and face information. Similar to the model creation process, if there are objects of non-mesh types in the file, BlenderPhotonics automatically converts these objects to mesh types.

### 2.5 Data exchange between Blender and Octave via portable JSON/JMesh files

The data exchange between Blender and Octave is achieved via the combination of Blender’s Python API `bpy`, the Octave-Python interface `oct2py`, JSON based data exchange Python module `jdata` and the built-in JSON parser in Iso2Mesh. Although the default data exchange scheme in `oct2py` is achieved using MATLAB’s proprietary .mat format, we intentionally adopted the JSON-compatible JMesh format^42^ (.jmsh) because of the following advantages: 1) JMesh is human-readable while MATLAB .mat file is not, 2) JMesh is a plain JSON file that can be readily read/written in nearly all programming environments with lightweight parsers (many are built-in to the programming languages, such as Python, Perl, MATLAB and JavaScript), 3) JMesh is directly editable and can be version-controlled while .mat files are binary-only, 4) last but not the least, JMesh/JSON files can be readily used for web applications or hierarchical databases (such as NoSQL databases) while .mat files require specialized parsers that are not widely available. We want to highlight that JMesh is not a new format, but rather a set of JSON-compatible “name”/”value” pairs representing mesh-based data structures. Existing mesh data stored in other formats, such as Visualization Toolkit (VTK) format,^43^ can be potentially converted to JSON/JMesh and used in BlenderPhotonics via Octave-based parsers or external converters.

In Fig. 4, we use a simple tetrahedral mesh (of a unit cube) containing 8 nodes, 12 surface triangles and 6 tetrahedral elements, shown in Fig. 4(a)) as an example to illustrate the simplicity of using JSON/JMesh data annotations to store mesh data. Because the JMesh format is built upon the JData specification^38, 44^ – a lightweight standard using JSON to annotate scientific data, JMesh mesh data constructs also supports strongly-typed array (see Fig. 4c) and record-level binary data compression^44^ (see Fig. 4d). We strongly encourage developers of other biophotonics tools considering adopting such format in their applications to achieve easy interoperability between multiple tools and ease in future extensions.

**Fig 4.**
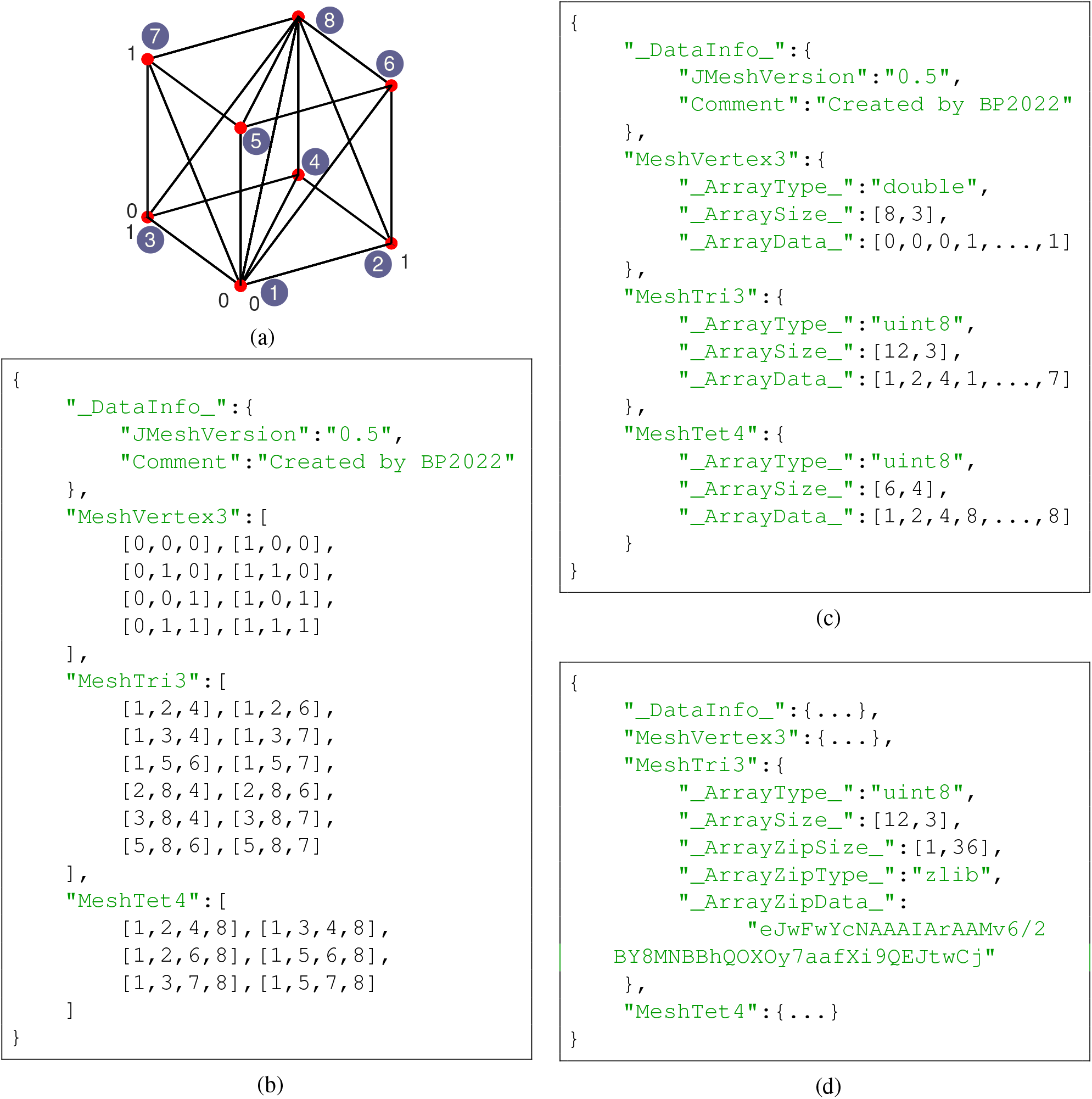
Sample JMesh representations of (a) a unit-cube (node numbering shown in circles). A JMesh file written with (b) plain JSON arrays, (c) JData-based annotated array and (d) compressed binary array are supported and parsed by our Python and MATLAB toolboxes. Note that JMesh node indices (such as in *“MeshTri3”* and *“MeshTet4”* constructs) start from 1.

### 2.6 Tetrahedral mesh generation from Blender scenes

The surface mesh models created in Blender, consisting of a single or multiple objects, are used as the input for the next step – tetrahedral mesh generation. The mesh generation process consists of two parts: surface mesh pre-processing and tetrahedral mesh generation. The pre-processing of the surface mesh is achieved in Blender. First Blender converts all non-mesh type objects to mesh types to get well-defined points and faces. Other types of objects in the scene that cannot be converted into a mesh object, such as light sources and cameras, will be excluded. All remaining objects are merged into one surface mesh object and checked for the presence of self-intersecting triangles. When self-intersection exists, Blender automatically inserts new points on the intersection line and split the faces, resulting in multiple sub-regions. At the end of the pre-processing step, a watertight non-self-intersecting triangular surface mesh is generated.

The pre-processed surface data are subsequently sent to Octave for mesh generation. Iso2Mesh is used to complete this step. First, the vertex coordinates and face information of the pre-processed model are saved as a .jmsh file and then loaded to Octave. In addition, several meshing parameters are prompt in a dialog (see Fig. 3) to the user to adjust the density of the output mesh, including 1) maximum tetrahedral element volume,^37^ and 2) percentage of surface mesh edges being kept after simplification, as defined in Iso2Mesh.^37^ The former parameter is passed to a mesh simplification algorithm implemented in the CGAL library^29, 30^ based on the Lindstrom-Turk algorithm.^45^ The latter parameter is passed to Tetgen to set the maximum volume of the generated tetrahedral mesh. The two parameters act together to control the density of the generated mesh. In Octave, the inputs for Iso2Mesh are node coordinates, face data, and mesh generation parameters. In addition,

Iso2Mesh calls Tetgen (version 1.5) for mesh generation and region labeling. After tetrahedral mesh generation, Iso2Mesh returns three output data arrays, namely, “node”, “face”, “elem” (see Fig. 5). They correspond to the vertices of the tetrahedral mesh, the faces triangle node indices, and the vertex indices of each tetrahedron, along with a label denoting the region where it belongs to, respectively. The tetrahedral mesh data are cached in a separate .jmsh file for subsequent calls. In addition, the exterior triangular surface for each tissue region/label is extracted and stored in a regional mesh .jmsh file to be loaded in Blender.

**Fig 5.**
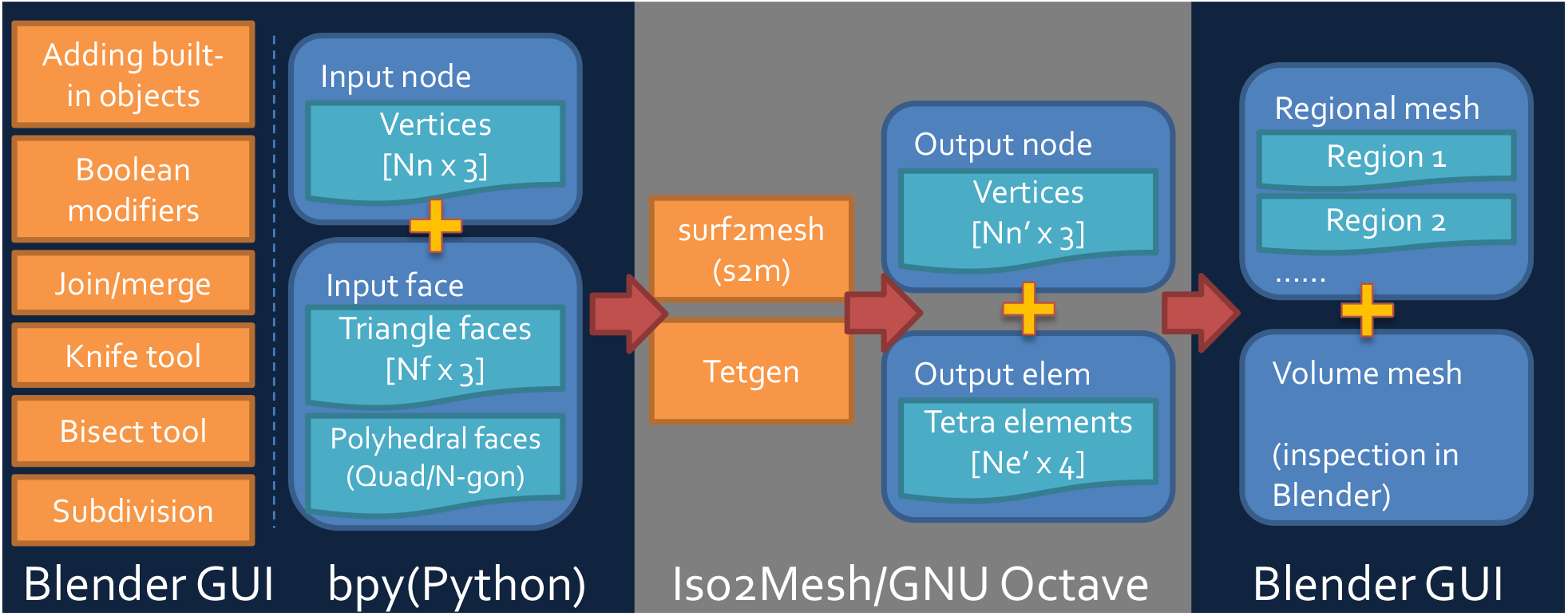
Mesh data structure exchanged between Blender and Iso2Mesh.

The successful generation of a tetrahedral mesh readily enables a user to perform an array of advanced modeling tasks, such as performing finite-element (FE) or boundary-element (BE) based analyses to solve the forward or inverse problems in the domains such as mechanics, computational electromagnetics (CEM), computational optics, computational fluid dynamics (CFD) etc. The created volume and surface meshes can be exported from Blender in the format that is acceptable by specialized computational physics or multi-physics solvers such as ANSYS, ABACUS, COMSOL etc. Even if a user does not have the need to perform subsequent numerical analyses, Blender/BlenderPhotonics offers efficient visualization, manipulation, and advanced transformation to a surface or tetrahedral mesh, making rendering and processing complex 3-D data with ease.

### 2.7 Tetrahedral mesh generation from 3-D volumetric images

For researchers in the fields of medical imaging and biophotonics, creating simulations from 3-D volumetric imaging data is a highly sought-after feature, but only limited tools are available for medical-image based 3-D mesh generation.^28^ Iso2Mesh is one of only a few tools^37^ that support this type of processing. In BlenderPhotonics, a dedicated submodule named “*Volume2Mesh*” is provided, as shown in Fig. 3. This module calls Iso2Mesh to automatically tessellate a 3-D volume stored in the NIfTI,^40^ JNIfTI^41^ or MATLAB .mat formats and output a surface mesh model for subsequent processing. NIfTI is an open-standard to store MRI/fMRI scans and is widely supported in the field of neuroimaging; JNIfTI is a standardized^41^ JSON wrapper^44^ to the NIfTI format to enable human-readability, easy extension and data-exchange between neuroimaging tools. By calling the streamlined mesh generation function in Iso2Mesh, the 3-D volume data from a JNIfTI/NIfTI file can be pre-processed to create both tetrahedral volume mesh as well as surface mesh that is at the outer surface of the volume mesh. Multi-labeled 3-D volume can also be directly processed by Iso2Mesh to create the surface mesh model. Each labeled sub-region is saved as a separate surface mesh record inside the exchange file. Unlike the previous two input types, the volume mesh and the regional mesh are generated simultaneously when a volume data file is imported into Blender. Nonetheless, a surface mesh extracted from the volume and regional mesh are displayed in Blender to allow manual editing and further manipulation in case additional complex features, such as growing hairs as shown in the later section, are desired.

To use this feature, one simply clicks on the file browser button under the “*Volume2Mesh*” section of BlenderPhotonics interface. This allows users to browse a text-JNIfTI (.jnii), binary-JNIfTI (.bnii), NIfTI (.nii/.nii.gz) or MATLAB .mat file stored in a local folder. In addition to loading a volume image from user’s local disk, one can also type in a URL pointing to an online JNIfTI/NIfTI/.mat file. BlenderPhotonics automatically downloads the online data file and reads the content. After either a valid file path or URL is supplied, one can click on the *“Convert 3-D image file to mesh”* button. A simple parameter dialog, shown at the middle of Fig. 3, is popped up, allowing users to set key meshing parameters, such as the upper-bound of tetrahedron volume (*V*_*max*_,^32, 37^ in cubic length unit), upper-bound of the Delaunay sphere radii of the surface triangles (*R*_*max*_,^29, 37^ in voxel unit), maximum allowed deviation (in voxel unit) from the voxelated boundary and mesh extract methods, including *“cgalmesh”*,^29^ *“cgasurf”*^30^ and *“simplify”*.^28^ Clicking on the *“OK”* button signals Iso2Mesh in Octave to start mesh generation. If successful, the regional mesh is loaded to Blender for inspection. If the mesh is not satisfactory, one can adjust the meshing setting and recreate the mesh.

### 2.8 Mesh-based Monte Carlo photon simulation workflow

In this work, we are particularly interested in solving the RTE inside complex media using the Monte Carlo (MC) method. Our previously developed MMC solver has already attracted a large and active user community consisting of students, researchers and academics from biomedical optics and optical neuroimaging domains.^23^ However, the lack of an intuitive simulation domain preparation interface makes it difficult to use for less-experienced users. With the development of BlenderPhotonics, we specifically address this issue by interfacing the 3-D mesh generation output from BlenderPhotonics with MMC and create a streamlined MC simulation environment in Blender. The MC simulation step requires three sub-steps: 1) domain preparation, 2) execution of photon simulation, and 3) output data visualization.

The domain preparation stage refers to the process of setting up the necessary optical simulation parameters, including optical properties for each tissue label, light source settings and global simulation settings such as total photon numbers etc. After the completion of the mesh generation step above, Blender/BlenderPhotonics loads and visualizes regional meshes while assigning four custom properties for each region, i.e. *µ*_*a*_ (1/mm), *µ*_*s*_ (1/mm), *g* and *n* (Fig. 6). At the same time, a light source is added in the 3-D domain. The light source has several user-customized properties such as 3-D position, orientation, source type, and total photon number to be simulated (Fig. 6). A user needs to set the optical parameters for each region of the mesh, place the light source in the desired position and adjust its orientation, and finally set the simulated photon number for the light source. It is important to note that the default unit of the domain is assumed to be mm in Blender-Photonics. Once a user completes these setup tasks, he/she can then start the optical simulation using a dedicated button on the BlenderPhotonics panel.

**Fig 6.**
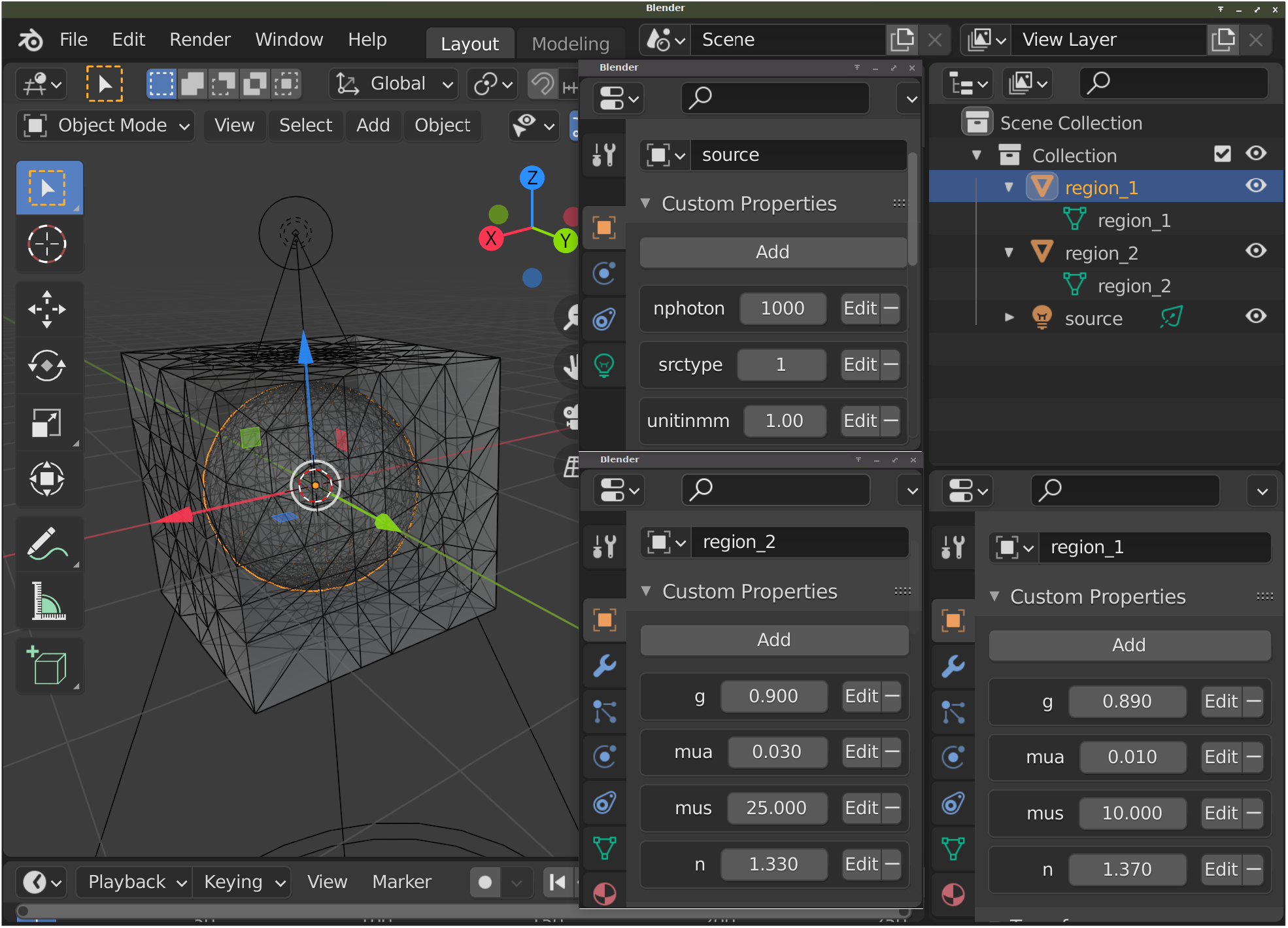
Optical parameters and light source configuration interface. The panels for setting optical parameters for region 1 (right) and region 2 (middle) are shown at the bottom; the panel for setting the source is shown at the middle-top.

When a photon simulation starts, BlenderPhotonics automatically reads the optical parameters of all region models in Blender, the light source information and passes it into Octave. Octave automatically generates a configuration file for the light simulation based on the supplied data. The mesh data of the simulation domain is derived from the mesh data saved during the tetrahedral generation stage. Other critical simulation parameters are passed from the user’s settings during the preparation stage. In particular, the light source direction is stored in Blender as a quaternion. In Octave, the direction in quaternions is used to compute the standard vector form.^46^ The total run-times of the MMC simulation depends on the model complexity, the total number of photons to be simulated and the optical properties. A progress bar, alongside with other simulation related information is printed in a command line window when MMC is being executed.

After MMC simulation is completed, the fluence map computed over the 3-D mesh is saved into a JMesh file and loaded back to Blender for visualization and post-processing. The fluence map is converted in the log-10 scale for better rendering of the simulation results. The log-scaled fluence intensity map are assigned as the “vertex weights” in Blender and rendered in pseudo-colors using a built-in color-map.

### 2.9 BlenderPhotonics file structure and optimization

BlenderPhotonics interacts with three types of files – Python scripts, Octave scripts and intermediate files. All Python scripts are placed in the add-on’s main directory. Octave scripts are stored in the `script` sub-folder inside the add-on’s directory. All intermediate outputs and data exchange files (JMesh files) are stored inside a temporary directory – for Linux/MacOS, this folder is typically `/tmp/iso2mesh-USER/blenderphotonics`, and for Windows, this is typically `C:\Users\USER\AppData\Local\Temp\iso2mesh-USER\blenderphotonics`, where *“USER”* should be replaced by the actual user account name.

In terms of run-times, BlenderPhotonics does not have direct impact to either those of Iso2Mesh mesh generation or MMC light simulation because those were performed in Octave. However, when writing and reading large numbers of surface mesh files, there is a noticeable data transfer overhead. Therefore, when loading the mesh data to Blender for inspection or rendering MMC simulation results, we only extract the surfaces of the regional mesh to reduce such overhead.

## 3 BlenderPhotonics Application Showcases

In this section, we first walk readers through three basic benchmarks to showcase the powerful yet easy-to-use processing pipleines of BlenderPhotonics – one for each of the 3 accepted input data types, including 1) “SkinVessel” benchmark for built-in shape based modeling, 2) “Colin27” benchmark for handling surface-mesh based inputs, and 3) “Digimouse” benchmark for 3-D volumetric data based modeling. For each benchmark, we first present an overview of the plug-in’s handling of the three models in operation and then report the screen captures following each processing step in BlenderPhotonics/Blender. Finally, we give readers two advanced examples to demonstrate the enormous possibilities to create complex and realistic tissue models using the rich arsenal of 3-D shape modeling tools offered by Blender. These two examples include 1) rough-surface modeling of skins, and 2) modeling complex human hairs, and understand how these complex realistic tissue features impact optical measurements.

### 3.1 “SkinVessel” benchmark – creating simulations from Blender objects

In the first example, we show mesh generation and light simulation of multi-layered skin tissues with an embedded blood vessel, adapted from the “SkinVessel” benchmark initially created by Dr. Steven Jacques for his MC software `mcxyz`.^47^ The model consists of a multi-layered slab structure, derived from a combination of a cube, dissecting planes, and a blood vessel created from a cylinder object in Blender, see Fig. 7. From bottom to top, the layers are named “Low-slab”, “Mid-slab” and “High-slab” respectively. The geometric and optical parameters of each region are reported in Table 1. The construction process of such model is shown in Figs. 7(a)-7(c).

**Table 1.** Geometric and optical parameters of the “SkinVessel” benchmark.

| Regions | $[z_{min}, z_{max}]$ (mm) | Radius (mm) | $\mu_a$ (1/mm) | $\mu_s$ (1/mm) | $g$ | $n$ |
| --- | --- | --- | --- | --- | --- | --- |
| Low-slab | [0.00, 0.10] | - | 0 | 1 | 1 | 1.37 |
| Mid-slab | [0.10, 0.16] | - | 1.657 | 37.594 | 0.9 | 1.37 |
| High-slab | [0.16, 1.00] | - | 0.046 | 35.654 | 0.9 | 1.37 |
| Cylinder | 0.50 (center) | 0.1 | 23.054 | 9.398 | 0.9 | 1.37 |

**Fig 7.**
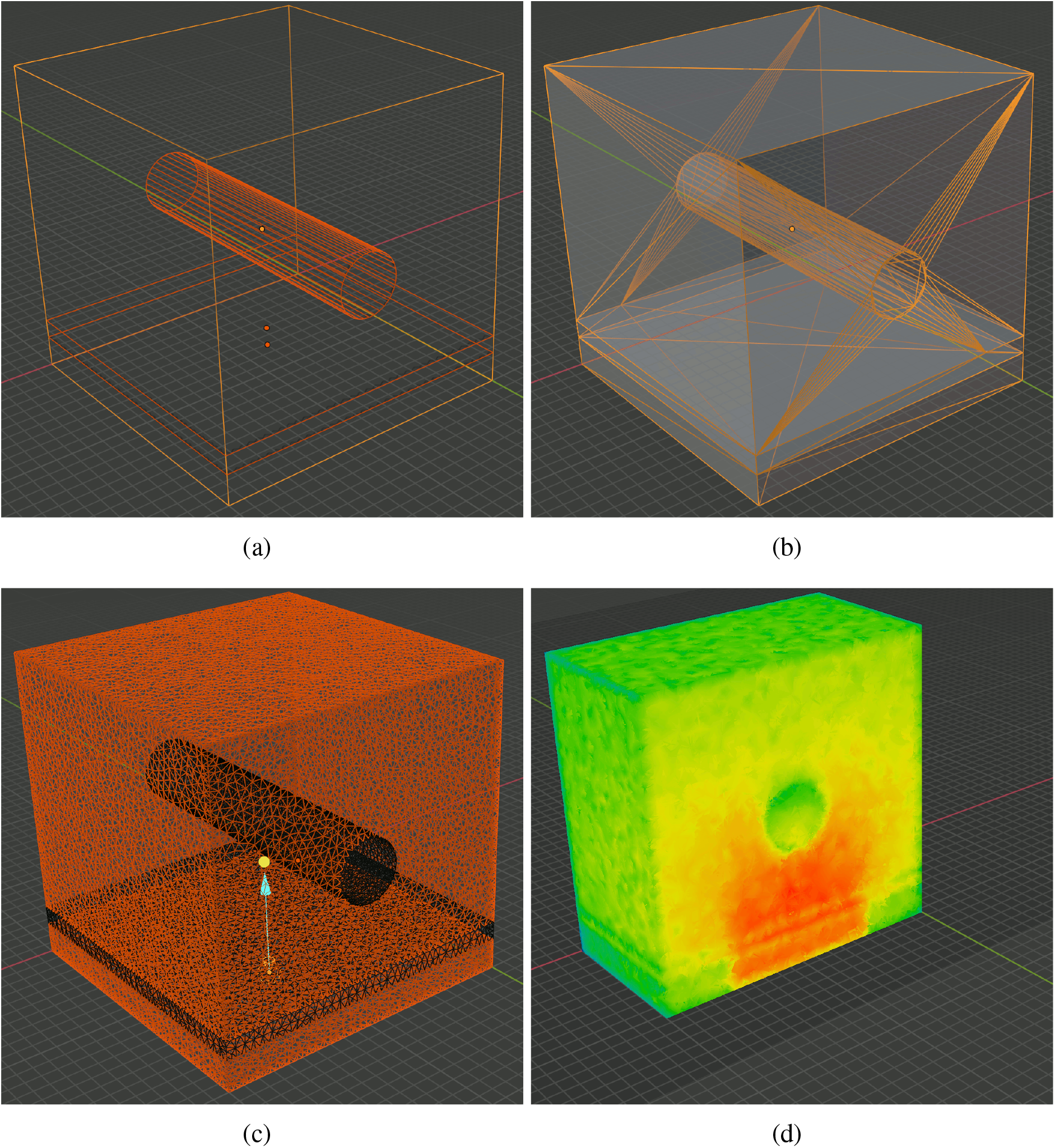
Intermediate steps of creating the SkinVessel benchmark: (a) after adding Blender objects, (b) after clicking “preview surface tesselation”, (c) output regional mesh and (d) fluence rate cross-section generated by MMC photon simulation.

To create this domain, we first use menu *“Add Mesh”* in the object-mode, to add a *“Cube”*, two *“Plane”* objects (Plane-1 and Plane-2), and a *“Cylinder”*. For each object, in the popup property dialog, we set their sizes (depth for the cylinder), offsets *x*_0_, *y*_0_, *z*_0_, rotation angles *ϕ*_*x*_, *ϕ*_*y*_, *ϕ*_*z*_ relative to each axis, and other properties according to Table 2. Here we set all objects lengths in voxel unit to match the original benchmark made for `mcxyz`; the voxel size (in mm) is specified by the *“unitinmm”* property shown in Table 2. This results in a 3-layered structure similar to Fig. 7(a). Once this model is created, one can click on the *“Preview surface tesselation”* button in BlenderPhotonics’s interface, and inspect the tessellated surface mesh, as shown in Fig. 7(b). One should see no self-intersecting triangles and each compartment of the surface must be watertight. After verification, select menu *“Edit\Undo”* to return to the polyhedral surface model. In the next step, one should click on the *“Convert scene to tetra mesh”* button on the BlenderPhotonics interface. In the shown dialog, we set the maximum tetrahedron volume to 30 (cubic length unit), optionally unchecked the *“Convert to triangular mesh”* while leaving other settings to default values. Clicking on *“OK”* allows Blender to call Iso2mesh and Tetgen to create a tetrahedral mesh of 109,110 nodes and 623,608 tetrahedra. Once completed, the volume mesh is loaded into the domain for inspection. This usually takes about 5-10 seconds.

**Table 2.** Blender properties set for each object to build the “SkinVessel” simulation. * this is achieved by setting the first element of the “srcparam1” property. All length units are in voxels to match the original `mcxyz` benchmark.

| Objects | Size/Depth | $x_0$ | $y_0$ | $z_0$ | $\phi_x$ | $\phi_y$ | $\phi_z$ | Other property settings |
| --- | --- | --- | --- | --- | --- | --- | --- | --- |
| Cube | 200 | 100 | 100 | 100 | 0 | 0 | 0 | - |
| Plane-1 | 200 | 100 | 100 | 20 | 0 | 0 | 0 | - |
| Plane-2 | 200 | 100 | 100 | 32 | 0 | 0 | 0 | - |
| Cylinder | 200 | 100 | 100 | 100 | 90° | 0 | 0 | radius=20, cap fill type=nothing |
| Source-1 | 50* | 100 | 100 | -10 | 180° | 0 | 0 | srctype=“disk”,unitinmm=0.005 |
| Source-2 | 50* | 100 | 50 | -50 | 150° | 0 | 0 | srctype=“disk”,unitinmm=0.0005 |

The next step is to configure an MMC simulation domain. To do so, one should click on the *“Load mesh and setup simulation”* button in the “*Multiphysics Simulations*” section of Blender-Photonics’s panel. This reloads the generated mesh in the previous step in the form of regional mesh (instead of a volumetric mesh), and name each tissue region as *“region i”* in the object list [Fig. 7(c)]. It also attaches default optical properties to each regional surface. In addition, a “light” object named *“source”* is also added to the scene positioned above the domain’s center. The custom property setting panels for each object is similar to those in Fig. 6. One should set the optical and source properties following Table 1. In this example, we use the settings for “Source-1”. Once completed, the domain is ready for the next step – MMC simulation – which is achieved conveniently by a single click on the *“Run MMC photon simulating”* button. In the popup dialog, one can adjust the simulation setting, such as maximum time-gate, reflection settings, normalization or choosing different GPU devices according to MMC’s command line options. Clicking on the *“OK”* button starts MMC simulation in the background. Once the simulation is completed, the computed light fluence map is loaded to the Blender rendering window and displays as “vertex weight” using a default color-map [Fig. 7(d)].

### 3.2 “Colin27” benchmark – creating simulations from surface meshes

In the second example, we demonstrate the steps needed to create tetrahedral mesh models and run MMC simulations using pre-created surface mesh models. The anatomical model was derived from a widely used human brain atlas, known as the “Colin27” atlas. A set of previously generated^23^ triangular surface meshes containing 4 closed tissue surfaces – scalp, ceroboralspinal-fluid (CSF), gray-matter (GM) and white-matter (WM) – are saved in the form of a JSON/JMesh file and loaded to Blender using the *“Import surface mesh”* button of BlenderPhotonics; a cropped input surface mesh can be found in Fig. 8(a). By a single click on the *“Convert scene to tetra mesh”* button in BlenderPhotonics and set 100 as the maximum element size, one can create a tetrahedral mesh with a cross-sectional plot shown in Fig. 8(b). Next, click on the *“Load mesh and setup simulation”* button, one can set the displayed regional meshes [Fig. 8(c)] using the optical parameters listed in Table 3, with the light source position, light source direction and photon number being [75.76, 66.99, 168.21] mm, [0.1636, 0.4569, −0.873] mm and 10^8^, respectively. Finally, a single click on the *“Run MMC photon simulation”* button starts MMC simulation using the input atlas and load the computed fluence map back to Blender once the simulation is completed [see Fig. 8(d)].

**Table 3.**
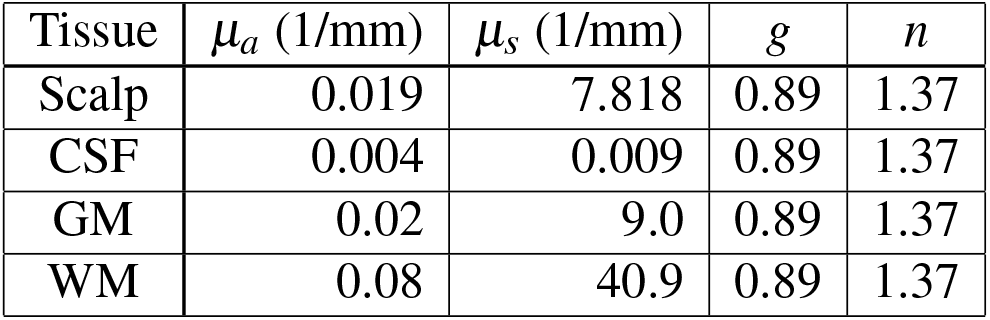
Optical properties for each tissue type in the Colin27 benchmark.

| Tissue | $\mu_a$ (1/mm) | $\mu_s$ (1/mm) | $g$ | $n$ |
| --- | --- | --- | --- | --- |
| Scalp | 0.019 | 7.818 | 0.89 | 1.37 |
| CSF | 0.004 | 0.009 | 0.89 | 1.37 |
| GM | 0.02 | 9.0 | 0.89 | 1.37 |
| WM | 0.08 | 40.9 | 0.89 | 1.37 |

**Fig 8.**
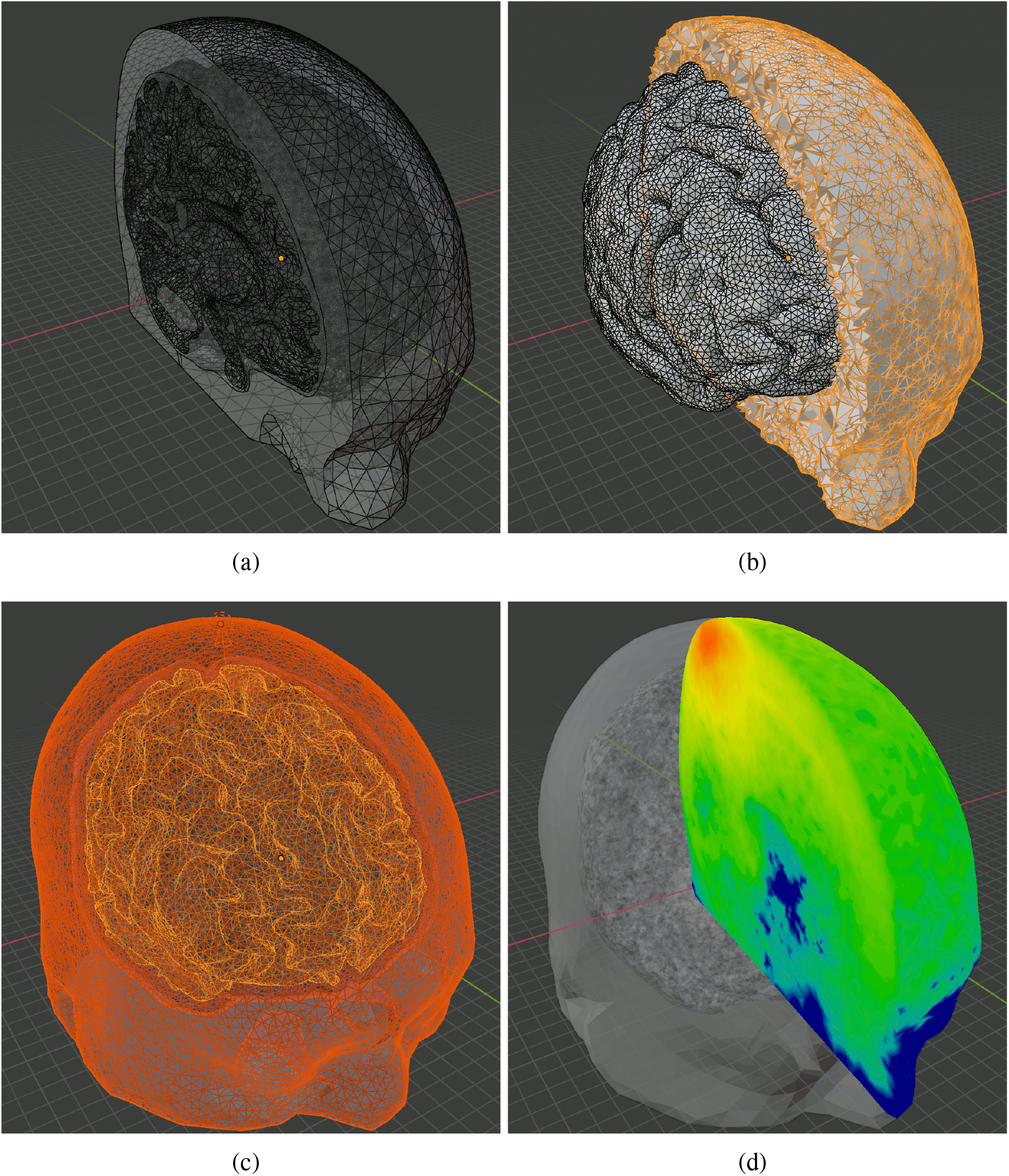
Intermediate steps of creating the Colin27 benchmark: (a) cropped input surface mesh, (b) cropped tetrahedral mesh, (c) region mesh and source, and (d) cropped fluence overlaid on the regional mesh.

### 3.3 “Digimouse” benchmark – creating simulations from segmented volumetric images

In this example, we show the processing steps converting a 3-D volume into a mesh model and subsequently run MC simulation. The benchmark is derived from a public dataset known as the “Digimouse” atlas, a segmented CT image of size 190 *×*496 *×*104 voxels with an isotropic voxel size of 0.8 mm. The original CT images of Digimouse atlas are converted to a JSON/JNIfTI file. To reproduce this example, one can directly type the file link “http://mcx.space/bp/data/digimouse. jnii“ in the JNIfTI file field in BlenderPhotonics’s interface. One can also download this file and browse it in the local disk. Next, one should click on the *“Convert 3-D image file to mesh”* button, set the maximum element volume to 100, and the volume image is loaded and processed in Iso2Mesh to create a tetrahedral mesh^12^ made of 21 tissue types [Fig. 9(b)]. The regional mesh for each tissue type [Fig. 9(a)] is then loaded back to Blender and the optical parameters of each region are assigned manually in Blender property dialog, according to those listed in Table 4. The light source position, light source direction and photon number are set to [40, 160, 80] mm, [0, 0, −1] mm and 10^8^ respectively. One should also set the *“unitinmm”* property of the source to the input voxel size, which is 0.8 (mm) in this case. The tetrahedral mesh of the atlas consists of 72,815 vertices and 407,739 tetrahedra. The final MMC photon simulation results are shown in Figs. 9(c)-9(d). To demonstrate the advanced rendering capability of Blender, we show two ways of creating cross-sectional images of the output – in Fig. 9(c), we can box-select a region of faces in the edit-mode and press “delete” on the keyboard. This creates a trimmed mesh showing the internal surfaces with the associated vertex-weight (fluence values). However, to create a flat cross-section, one can select *“Mesh\Bisect”* in the edit-mode, and draw a straight-line across the mesh. In the *“Bisect”* setting dialog, one can choose to remove either side of the bisecting plane, and fill the cross-sectional plane with flat patches. The result is shown in Fig. 9(d). A even faster way to crop a mesh is to use the view-port cropping feature by pressing shortcut *“Alt+B”* and box-select the region to be kept.

**Table 4.** Optical parameters for each of the regions/labels of the “Digimouse” benchmark.^48,49^

| Region ID | Tissue Type | $\mu_a$ (1/mm) | $\mu_s$ (1/mm) | $g$ | $n$ |
| --- | --- | --- | --- | --- | --- |
| 1 | skin | 0.0191 | 6.6 | 0.9 | 1.37 |
| 2 | skeleton | 0.0136 | 8.6 | 0.9 | 1.37 |
| 3 | eye | 0.0026 | 0.01 | 0.9 | 1.37 |
| 4 | medulla | 0.0186 | 11.1 | 0.9 | 1.37 |
| 5 | cerebellum | 0.0186 | 11.1 | 0.9 | 1.37 |
| 6 | olfactory bulbs | 0.0186 | 11.1 | 0.9 | 1.37 |
| 7 | external cerebrum | 0.0186 | 11.1 | 0.9 | 1.37 |
| 8 | striatum | 0.0186 | 11.1 | 0.9 | 1.37 |
| 9 | heart | 0.024 | 8.9 | 0.9 | 1.37 |
| 10 | rest of the brain | 0.0026 | 0.01 | 0.9 | 1.37 |
| 11 | masseter muscles | 0.024 | 8.9 | 0.9 | 1.37 |
| 12 | lachrymal glands | 0.024 | 8.9 | 0.9 | 1.37 |
| 13 | bladder | 0.024 | 8.9 | 0.9 | 1.37 |
| 14 | testis | 0.024 | 8.9 | 0.9 | 1.37 |
| 15 | stomach | 0.024 | 8.9 | 0.9 | 1.37 |
| 16 | spleen | 0.072 | 5.6 | 0.9 | 1.37 |
| 17 | pancreas | 0.072 | 5.6 | 0.9 | 1.37 |
| 18 | liver | 0.072 | 5.6 | 0.9 | 1.37 |
| 19 | kidneys | 0.05 | 5.4 | 0.9 | 1.37 |
| 20 | adrenal glands | 0.024 | 8.9 | 0.9 | 1.37 |
| 21 | lungs | 0.076 | 10.9 | 0.9 | 1.37 |

**Fig 9.**
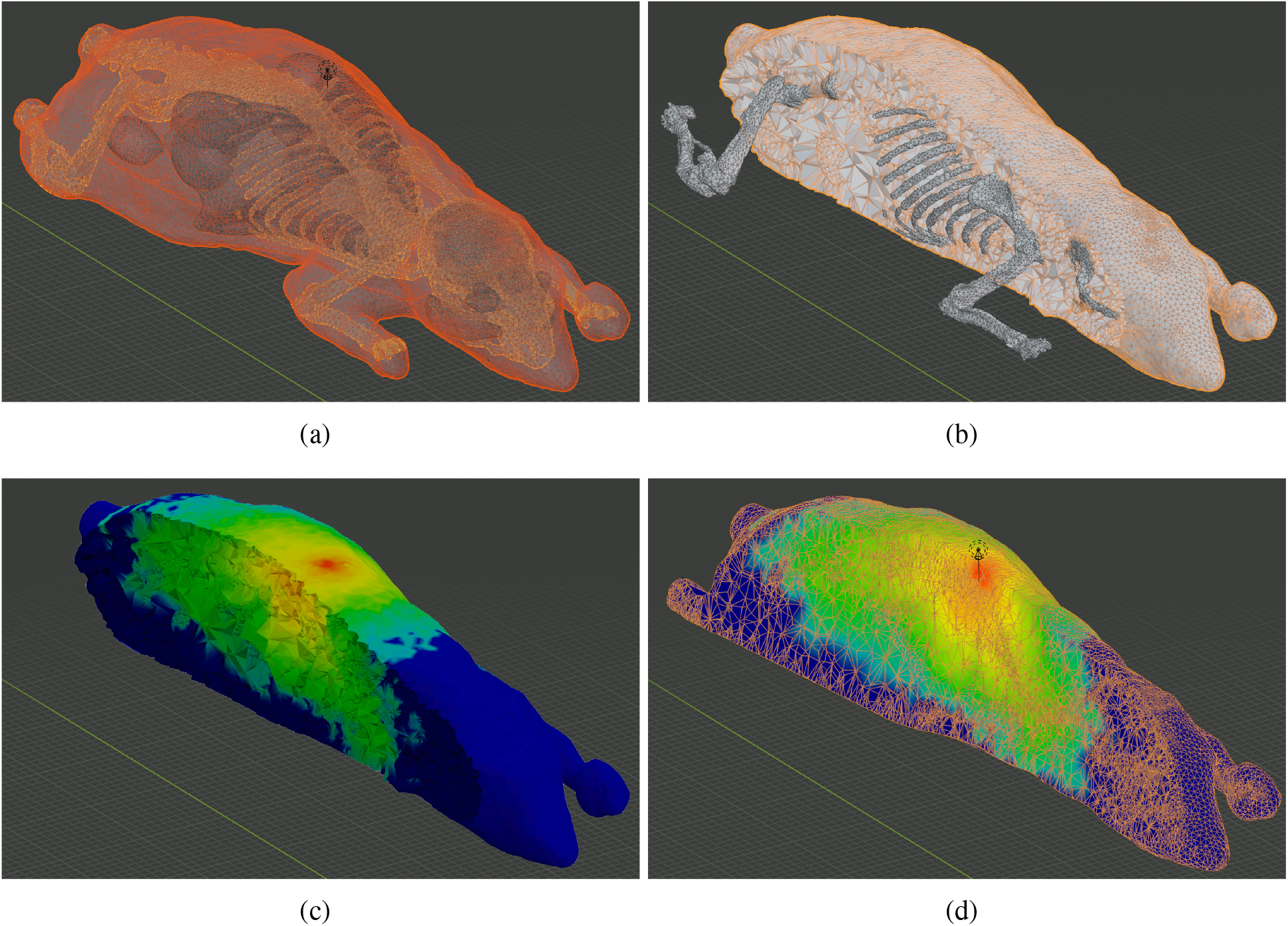
Intermediate steps of creating the Digimouse benchmark: (a) regional mesh after converting the volume to mesh, (b) output tetrahedral mesh (cropped), (c) fluence-rate map rendering over a trimmed mesh using face-deletion and (d) that rendered over a sliced mesh using the “bisect” function.

### 3.4 Advanced modeling example 1 – simulating rough surfaces using BlenderPhotonics

BlenderPhotonics’ ability to bridge Blender’s superior model creation/transformation capability with state-of-the-art 3-D mesh generation and quantitative photon simulation opens numerous possibilities to advance our understandings to complex tissue-photon iterations in a realistic setting, helping us design better instrument and experiments and accounting for scenarios that simple models could never reveal. We can not enumerate all possible advanced modeling functions provided by Blender, however, just to provide inspirations for biomedical optics researchers and computational scientists, we hand-picked two features that are relevant to tissue optics. In the first example, we use the rough-surface creation feature in Blender and investigate how changing the roughness of a skin-mimicking surface could lead to different light distributions and results. In the second example, we can combine BlenderPhotonics with one of our most recent advances in MC simulation – the implicit MMC (iMMC) algorithm^27^ – to potentially enable the study of the impact of human hairs in fNIRS measurements. To the best of our knowledge, these types of studies have not been reported by other MC simulators.

In the rough-surface simulations, we adjusted the geometric parameters of the interface based the aforementioned “SkinVessel” benchmark to simulate the roughness of realistic skin surface, which was difficult to achieve in the past due to the lack of relevant features in the mesh generation tool. In this case, all parameters used in this benchmark are largely the same as we described in the previous subsection, except that we change the source position from those for “Source-1” to those of “Source-2” according to Table 2. We also shrink the domain by a factor of 10 to amplify the effects by changing the length unit, defined as a custom property named *“unitinmm”* attached to the source object, from 0.005 (mm) to 0.0005 (mm). In addition, we modified the Plane-1 object (see Table 1) to create a rough surface. This is achieved by first selecting the plane object in the edit-mode, right-clicking on the object, choosing *“Subdivide”*, and setting the division number to 39 [Fig. 10(a)]. In the next step, one should switch to the vertex-selection mode, and select menu *“Mesh\Transform\Randomize”*, then set *“Amount”* to 1, and *“Normal”* to 1.0. This creates a maximum *±*1 *×* 0.0005 mm normal-direction-constrained random movement [Fig. 10(b)]. According to literature,^50^ the roughness of the human skin surface can be described using the parameter *R*_*a*_, defined as the arithmetic mean deviation of the depth profile:

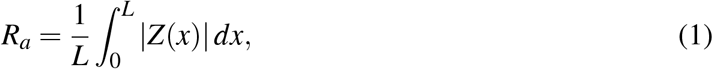

where *Z*(*x*) is the deviation from the average depth at *x* and *L* is sampling length. In this case, the *R*_*a*_ value of the surface is 0.25 *µ*m [Fig. 10(b)] and 1.5 *µ*m [Fig. 10(c)] in two separation simulations. The optical parameters for each domain are given in the Table 2. To test the effect of light transmission across a rough-surface, the light source was set as a disc-like light source with a radius of 25 *µm*, positioned off-center 30 degree tiled towards the +*x*-axis (i.e. “Source-2” in Table 2). The results of the fluence distributions for *R*_*a*_ = 1.5*µ*m are shown in Fig. 10(f). In comparison, the light simulation for a low roughness skin at *R*_*a*_ = 0.25*µ*m is shown in Fig. 10(e).

**Fig 10.**
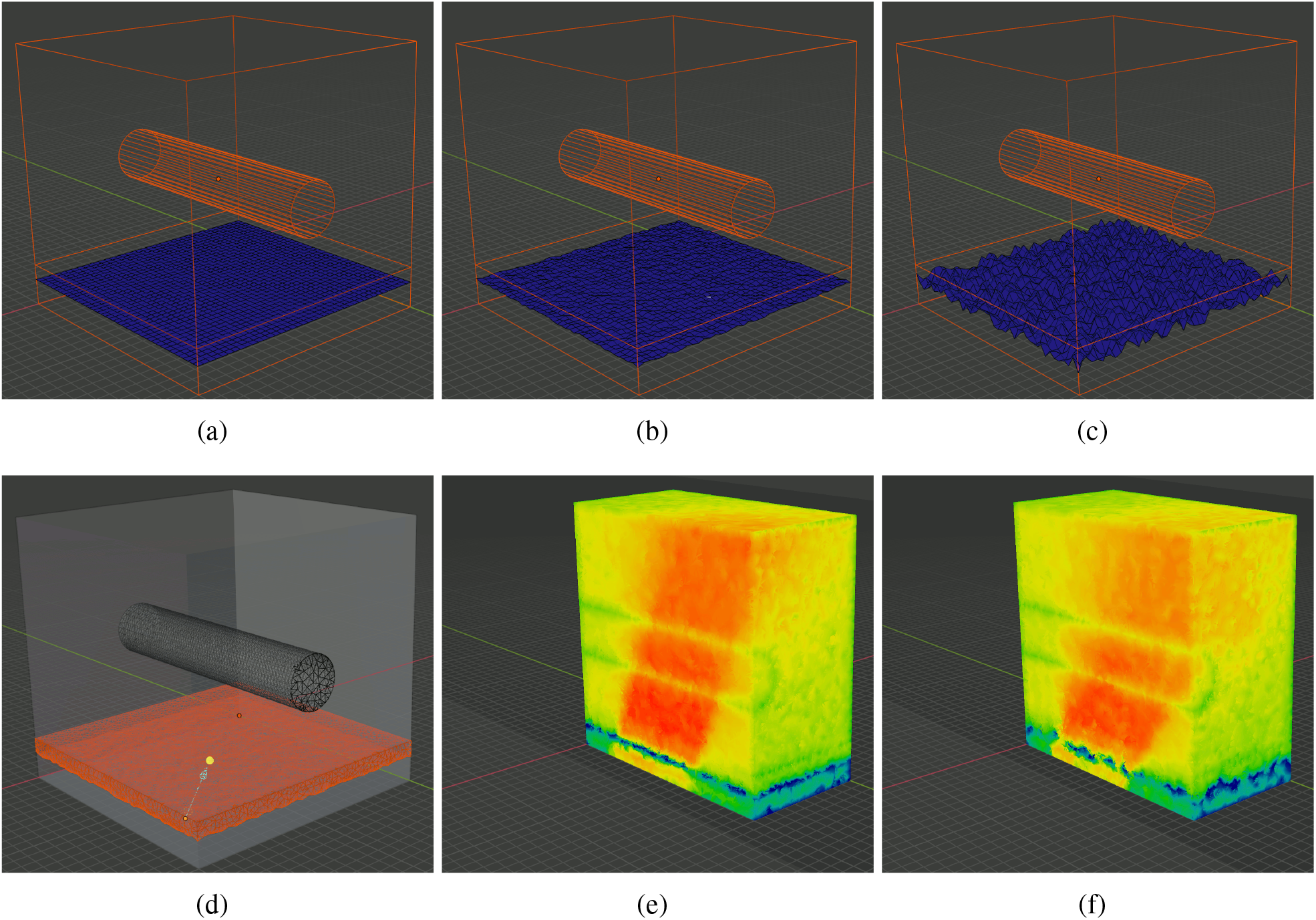
Comparisons between (b,e) low-roughness and (c,f) high-roughness surfaces using BlenderPhotonics: (a) initial surface model after subdivision, (b,c) created rough surfaces using randomized nodal offsets, and (e,f) MMC simulated fluence rate cross-sectional plots from a tilted disk-source.

The images shown in Figs. 10(e)-10(f) demonstrates that considering tissue surface roughness can noticeably alter the incident light direction. This is because a rough-surface acts as a diffuser and broadens the incident beam, significantly reducing the collimation of the incident beam. The capability of BlenderPhotonics to simulate such realistic surface model can assist researchers to better optimize their imaging system and obtain more accurate quantification.

### 3.5 Advanced modeling example 2 – simulating realistic human hairs using Blender hair system

In addition to the roughness of realistic human skin, hair is also widely considered an important factor when performing optical measurements on the human body. In particular, the presence of human hair is considered one of the most challenging aspects of fNIRS measurements.^51^ Unfortunately, due to the high complexity of modeling realistic hair, previous publications on hair modeling were limited to simple models involving only dozens of hairs of uniform hair-root distribution and tilting angle. Here, we combine the advanced hair modeling system in Blender with the latest advances in the implicit MMC (iMMC) simulation algorithm and provide a viable path for researchers to rigorously simulate the effects of hair in realistic measurements. A comprehensive study characterizing the impact of hair of different densities, colors, lengths, etc., in the context of fNIRS will be reported in a separate publication. Here, we just want to demonstrate the rich and advanced hair model generation capability provided by Blender and hopefully motivate readers to explore other complex shape modeling features provided by this open-source platform.

In this example, we first show results from “growing” realistic hairs over a simple 3-layered head model and then show hair models created using the aforementioned Colin27 atlas. First, we create a 3-layered slab model similar to the steps used in Section 3.1. The slab has dimensions of 20*×* 50 *×*34.51 mm^3^, with a 12.44 mm top layer simulating scalp/skull, a 2.07 mm middle-layer simulating CSF, and a 20 mm bottom layer simulating the brain (GM/WM). To grow hairs on the top surface, we first select the top surface in the edit-mode, right-click and select *“Subdivision”*, with number of cuts set to 1. Then select the face-center-vertex inserted by the *“Subdivision”* step and open the *“Object properties”* panel on the right. Under the *“Vertex group”*, add a new group, and click *“Assign”* and *“Select”* buttons in order to apply particle simulations over the selected surface later. Next, one should switch to the *“Particle properties”* panel, click on the plus-sign to create a new particle property, and then select *“Hair”*. This setting immediately grows 1000 straight (along the normal direction) hairs with random roots on the entire surface of the slab. To constrain the hairs to only the top-surface, go to the *“Vertex group”* sub-panel, under the particle properties, and choose the vertex group created earlier. Once a user switches to the object-model, the hairs should be displayed similar to Fig. 11(a). One can use the *“Particle properties”* panel to adjust various properties of the hair model, such as total count, length, tilting angle, randomness of the tilting angle, etc. For example, setting the *x/y/z* velocity under the *“Velocity”* panel changes the direction of the hairs and randomness of the tilting angle; setting hairs to tilt 45-degree in the +*x*-axis with a randomness of 0.2 results in a hair model shown in Fig. 11(b). To create even more realistic hair shapes, one can check the *“Hair Dynamics”* checkbox, and click the *“Play”* button at the bottom of the window. This applies forces to each segment of the hair shaft and creates curved hairs pulled by gravity. Dynamically curved hairs using Fig. 11(b) as the initial model results in Fig. 11(c). Similarly, this process can also be repeated in arbitrarily complex shapes, such as the Colin27 atlas. In Figs. 11(d)-11(e), we show the outcome of growing 80,000 hairs (165 hairs/cm^2^) with and without hair dynamics, respectively, on the Colin27 head surface; in Fig. 11(f), we increase the hair count to 400,000 (825 hairs/cm^2^) with hair dynamics enabled to simulate extremely dense and realistic human hair. A special script is used to export all hair vertices and the head mesh into a file that is processed by the Iso2Mesh toolbox to create iMMC simulation models for subsequent light transport simulations.^27^

**Fig 11.**
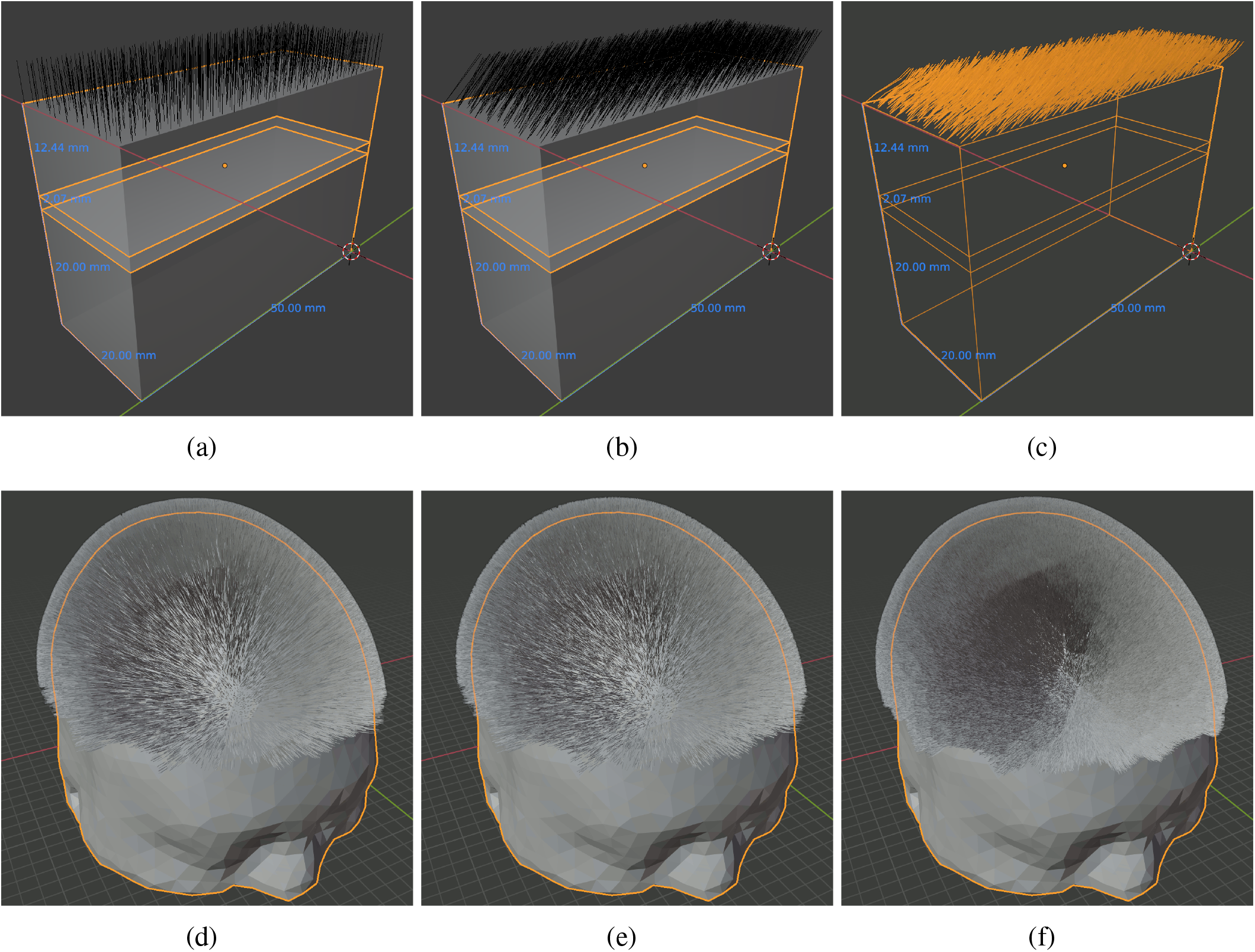
Demonstrations of different hair models created in Blender. In (a-c), we show (a) straight, (b) 45-degree tilted and (c) gravity-curved hairs grown over a 3-layered slab head model. We also show (d) straight and (e,f) gravity-curved hairs simulated over the Colin27 atlas at two densities: (d-e) 165 hairs/cm^2^ and (f) 825 hairs/cm^2^.

## 4 Conclusion

Despite the ample progress made towards developing open-source MC photon simulators, ease-of-use is still greatly lacking among most publicly available MC packages. This creates a barrier of entry for less experienced users and hampers the widespread use of these state-of-the-art simulation tools. The goal of this tutorial is to report on a novel software environment that utilizes the open-source modeling tool, Blender, and combines interactive 3-D mesh generation with streamlined MC light simulation, thereby making it possible to create sophisticated and life-like complex biophotonic simulations without needing to write a single line of code. With the BlenderPhotonics add-on developed in this work, one can create simulation domains interactively using the intuitive Blender interface while also taking advantage of the exquisite modeling toolkit offered by the Blender ecosystem and the associated large user community. This work greatly eases the utility of and expands the potential user community for MC simulation packages.

Using three basic examples, we show users the step-by-step handling of various types of inputs, including surface meshes, volumetric images, and Blender built-in objects, to conveniently create accurate and simulation-ready domain structures. Most of the examples only need a minute or two to create and only require a few clicks in Blender/BlenderPhotonics; such simulations can be saved, shared, and reproduced by other users. In addition, we demonstrate a significant expansion in our capability to create sophisticated and realistic simulations, enabled by BlenderPhotonics. In one of the advanced examples, we show detailed steps to create rough surfaces to simulate realistic human skin and demonstrate evidence that different roughness levels impact how light interacts with the underlying tissue.

Additionally, we provided a showcase demonstrating how to use BlenderPhotonics to create increasingly sophisticated human hair models. The hair/particle system is a well developed feature in Blender’s toolbox for building photo-realistic 3-D models. BlenderPhotonics makes the hair-generation function readily available for advanced fNIRS modeling. To this end, we have demonstrated the capability to control various hair features, such as length, density, root location, tilting angles, as well as gravity based bending.

In the next step, our focus is to further expand the BlenderPhotonics interface to incorporate additional existing features provided in Iso2Mesh and MMC, and make them accessible in an intuitive and interactive fashion inside Blender. In addition to MC simulations, we will also incorporate our Redbird^17, 52^ FEM DE forward/inverse solver with BlenderPhotonics. Moreover, we will continue developing the JMesh specification and enhance our current Blender-to-JSON export and import capability. Our goal is to establish JSON/JMesh/JNIfTI as the “source-code” format for scientific data, including mesh/shape and imaging related data. Converging towards a human-readable, universally supported, and easily extensible format is an important step towards enhancing interoperability between increasingly complex data analysis pipelines and reproducibility of both experiments and simulations in scientific research.

To conclude, BlenderPhotonics is an open-source Blender add-on, capitalizing upon the large and active Blender 3-D modeling ecosystem, that is able to serve as an interactive front-end for both 3-D mesh generation and 3-D mesh-based MC optical simulations. A simple yet functional GUI design enables users without prior coding experience to create complex tissue models, tessellate 3-D tetrahedral meshes, and, when needed, run streamlined MC photon simulations with ease. This software significantly shortens the learning curve for novice users of Iso2Mesh and MMC, allowing them to focus on data analysis rather than pre-processing. BlenderPhotonics, with its intuitive and feature-rich visual interface, is a great addition to the growing body of open-source MC simulators and helps disseminate these research software tools towards a broader research community. The latest BlenderPhotonics software can be downloaded at http://mcx.space/bp.

## Disclosures

No conflicts of interest, financial or otherwise, are declared by the authors.

## Acknowledgments

This research is supported by the National Institutes of Health (NIH) grants R01-GM114365, U24-NS124027, and R01-EB026998. The authors would like to thank Edward Xu for his help in preparing this manuscript.

**Yuxuan Zhang** is currently working toward the Ph.D. degree in Biomedical Engineering Department, University of Connecticut, Storrs, USA. He received his B.E. degree from Zhengzhou University, Zhengzhou, China, in 2019 and M.S. in Bioengineering from Northeastern University in 2021. His research interests include CRISPR-based biosensor, neuroelectronic interfaces and optogenetics, with a goal of developing advanced neural probes for modulation and monitoring of the nervous systems.

**Qianqian Fang**, PhD, is currently an Associate Professor in the Bioengineering Department, Northeastern University, Boston, USA. He received his PhD degree from Thayer School of Engineering, Dartmouth College, in 2005. He then joined Massachusetts General Hospital and became an Instructor of Radiology in 2009 and Assistant Professor of Radiology in 2012, before he joined Northeastern University in 2015. His research interests include translational medical imaging devices, multi-modal imaging, image reconstruction algorithms, and high performance computing tools to facilitate the development of next-generation imaging platforms.

## Notes

http://mcx.space/bp

https://github.com/COTILab/BlenderPhotonics

### Competing Interest Statement

The authors have declared no competing interest.

### Summary of Updates

- Describe possibilities of integrating with other MC simulators - additional comments regarding JMesh - updated title - removed installation script (old Fig. 3)

